## Supplementary Figures for "Arabidopsis TRB proteins function in H3K4me3 demethylation by recruiting JMJ14"

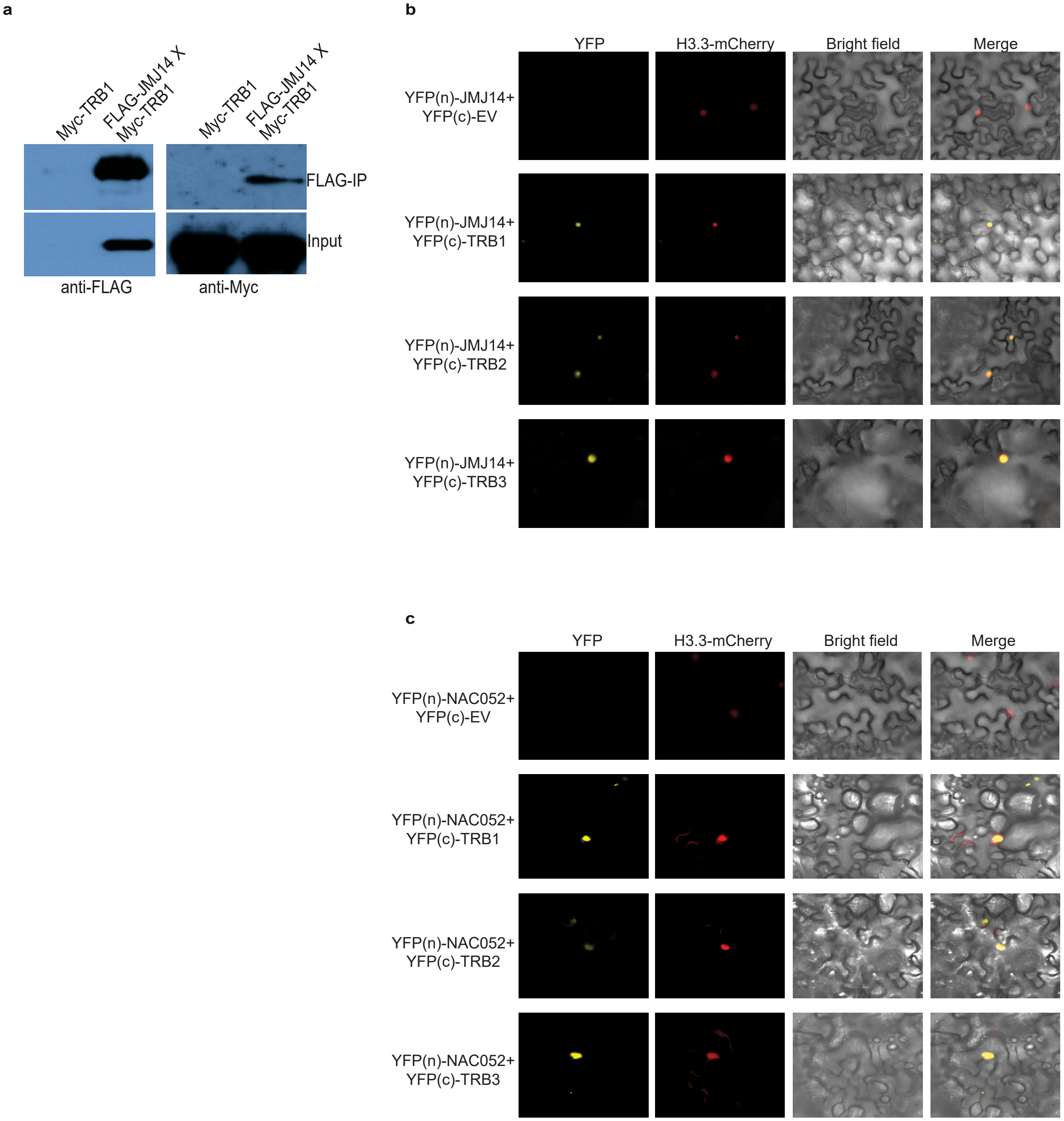


**Supplementary Fig. 1 FLAG-TRB1 interacts with Myc-JMJ14. a** Western blot showing the Co-immunoprecipitation (Co-IP) assay in FLAG-JMJ14 and Myc-TRB1 F2 crossed lines. **b-c** BiFC assays showing the YFP signals of leaves transiently co-expressing cYFP-TRBs and nYFP-JMJ14 (**b**) or nYFP-NAC052 (**c**).

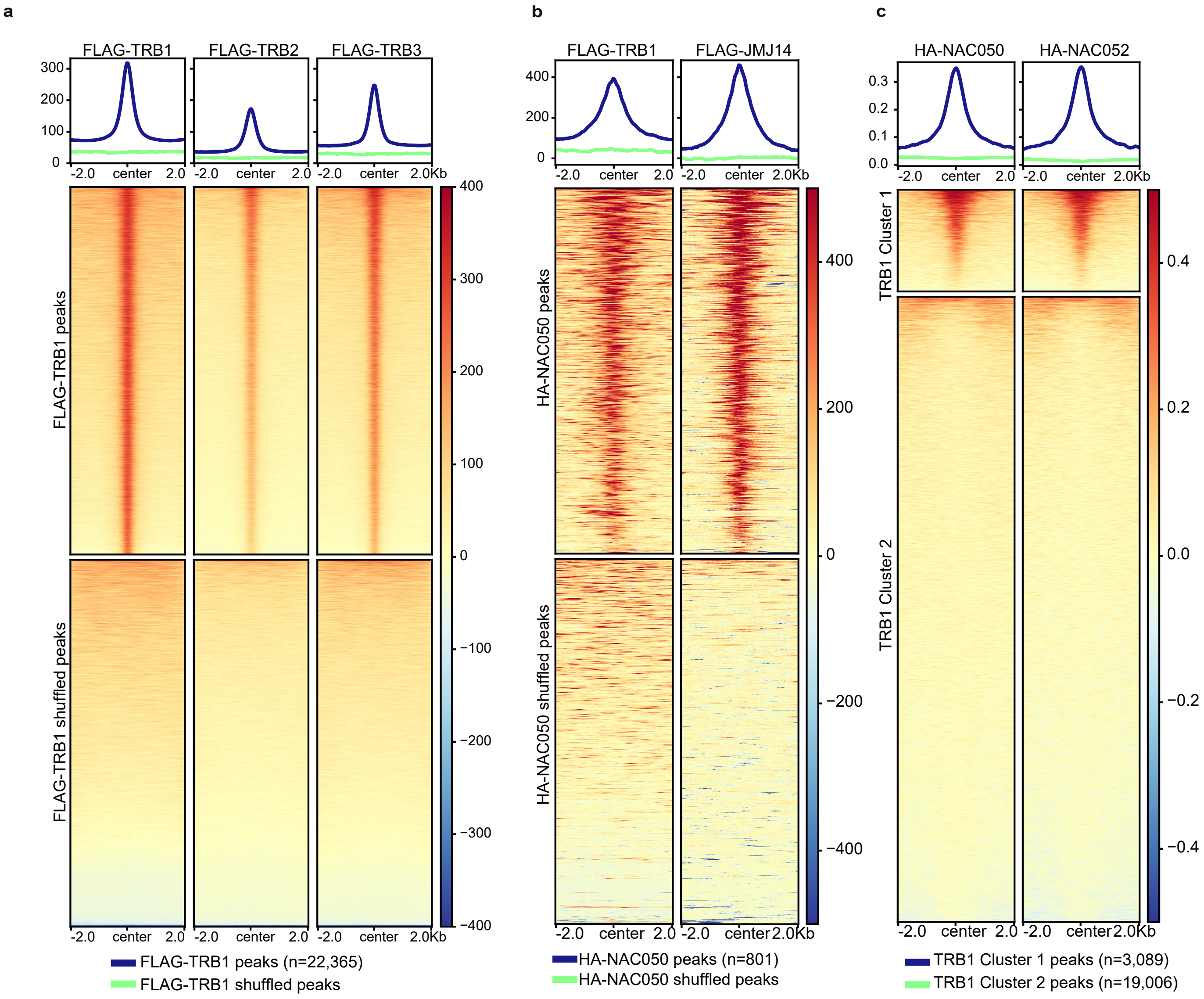


**Supplementary Fig. 2 TRBs are partially colocalized with JMJ14, NAC050, and NAC052. a-c** Metaplots and heatmaps showing FLAG-TRB1, 2, 3 ChIP-seq signals over FLAG-TRB1 peaks and shuffled peaks (**a**, n=22,365), FLAG-TRB1 and FLAG-JMJ14 ChIP-seq signals over HA-NAC050 peaks (n=801) and shuffled peaks (**b**), and HA-NAC050 and HA-NAC052 ChIP-seq signals over the TRB1 Cluster 1 peaks (n=3,089) and the Cluster 2 peaks (n=19,006) (**c**).

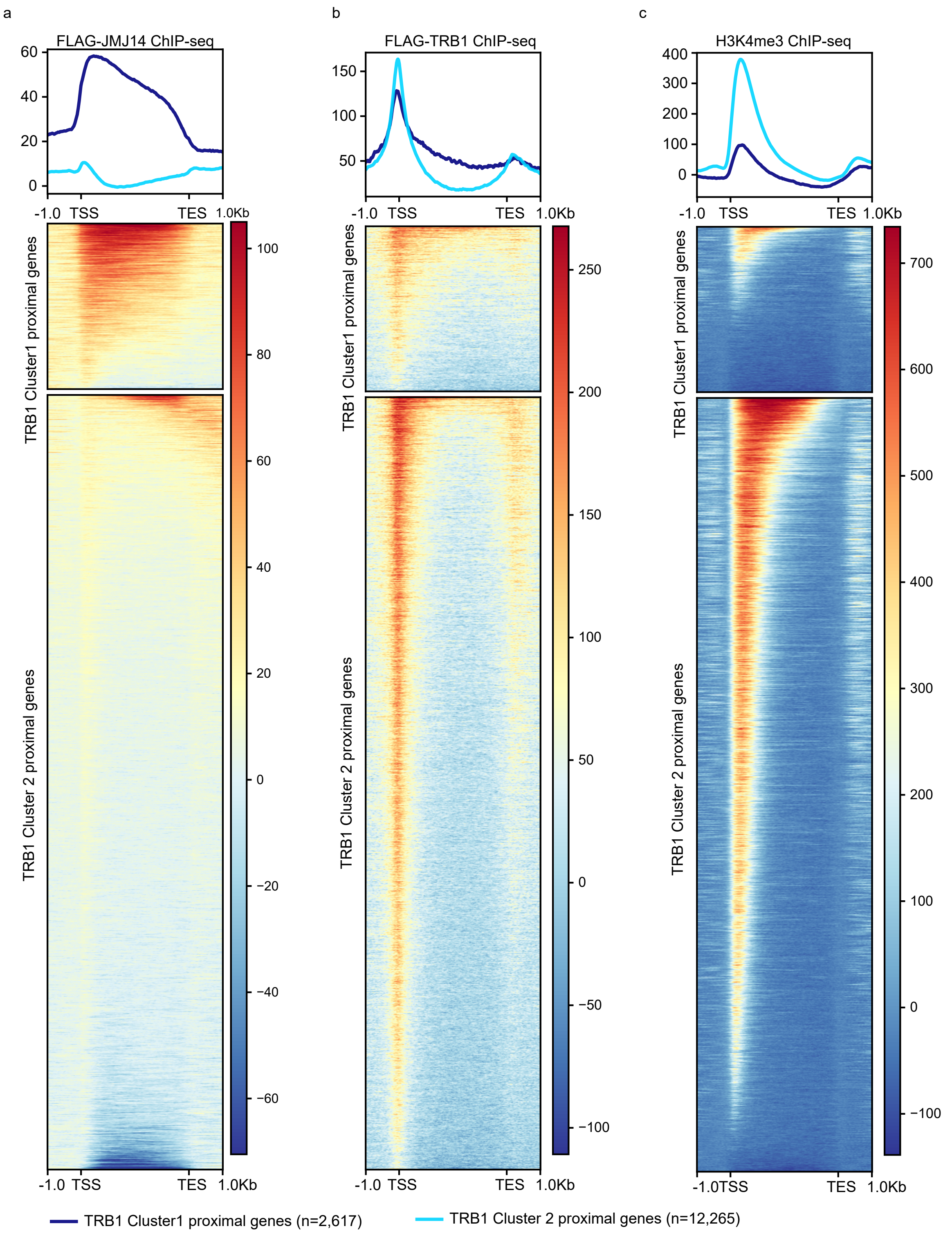


**Supplementary Fig. 3 JMJ14 and TRB1 are colocalized at gene body regions. a-c** Heatmaps and metaplots representing FLAG-JMJ14 (**a**), FLAG-TRB1 (**b**), and H3K4me3 (**c**) ChIP-seq signals over the TRB1 Cluster 1 proximal genes (n=2,617) and the Cluster 2 proximal genes (n=12,265).

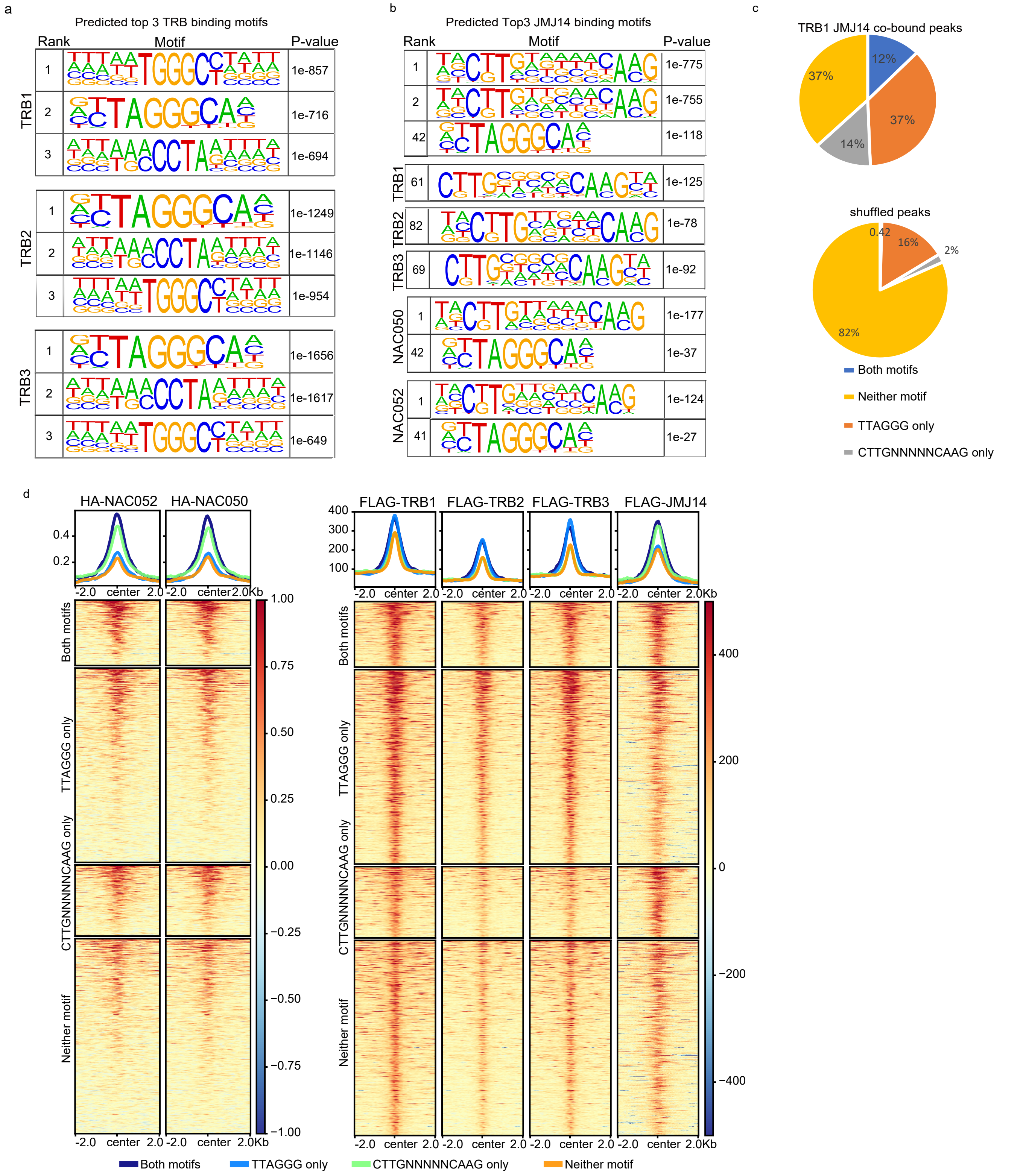


**Supplementary Fig. 4 Predicted TRB binding motifs include the telomeric repeat DNA sequence TTAGGG and the known JMJ14 binding motif CTTGnnnnnCAAG. a** Motif prediction by Homer showing the top 3 binding motifs of TRB1, TRB2, and TRB3. **b** Motif prediction by Homer showing that JMJ14, TRBs, NAC050, and NAC052 are all predicted to bind the TTAGGG and CTTGnnnnnCAAG motifs. **c** Pie charts indicate the percentile of the TRB1 and JMJ14 co-bound peaks (top panel) and shuffled peaks (bottom panel) displaying the TTAGGG motif only (n=1,176), the CTTGnnnnnCAAG motif only (n=431), both motifs (n=396), and neither (n=1,180), respectively . **d** Heatmaps and metaplots representing the normalized HA-NAC050/052, FLAG-TRB1, FLAG-TRB2, FLAG-TRB3, and FLAG-JMJ14 ChIP-seq signals over TRB1 and JMJ14 co-bound regions displaying the TTAGGG motif only, the CTTGnnnnnCAAG motif only, both motifs, and neither, respectively.

*trb2/trb3* Col-0

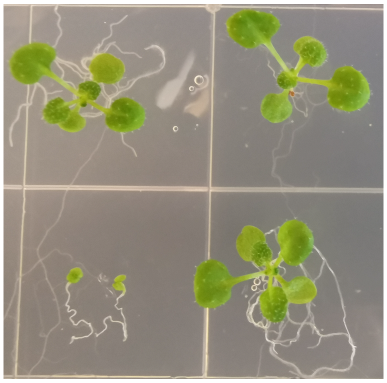

*trb1/2/3* *trb2/trb3 homo- trb1heterozygous*

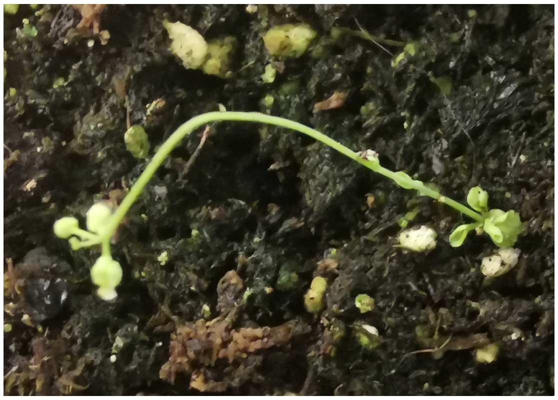

*trb1/2/3*

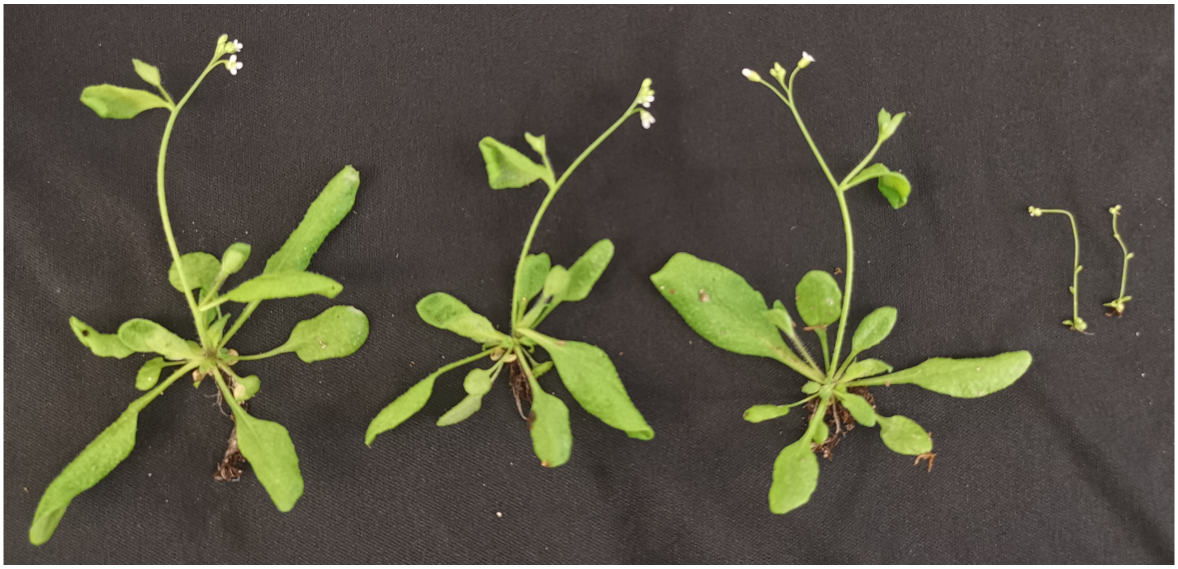

Col-0 *trb2/trb3* *trb2/trb3 homo- trb1 heterozygous* *trb1/2/3*

**Supplementary Fig. 5 The *trb1/2/3* triple mutant shows strong morphological defects.** The photos show the phenotype of *trb2/3* double mutant, Col-0, *trb1/2/3* triple mutant, and *trb2/3* homo- *trb1* heterozygous mutant grown on MS medium for 2-3 weeks (top left), and then transferred to soil for another 2 weeks (bottom). The top right photo shows a closer view of a *trb1/2/3* triple mutant on soil.

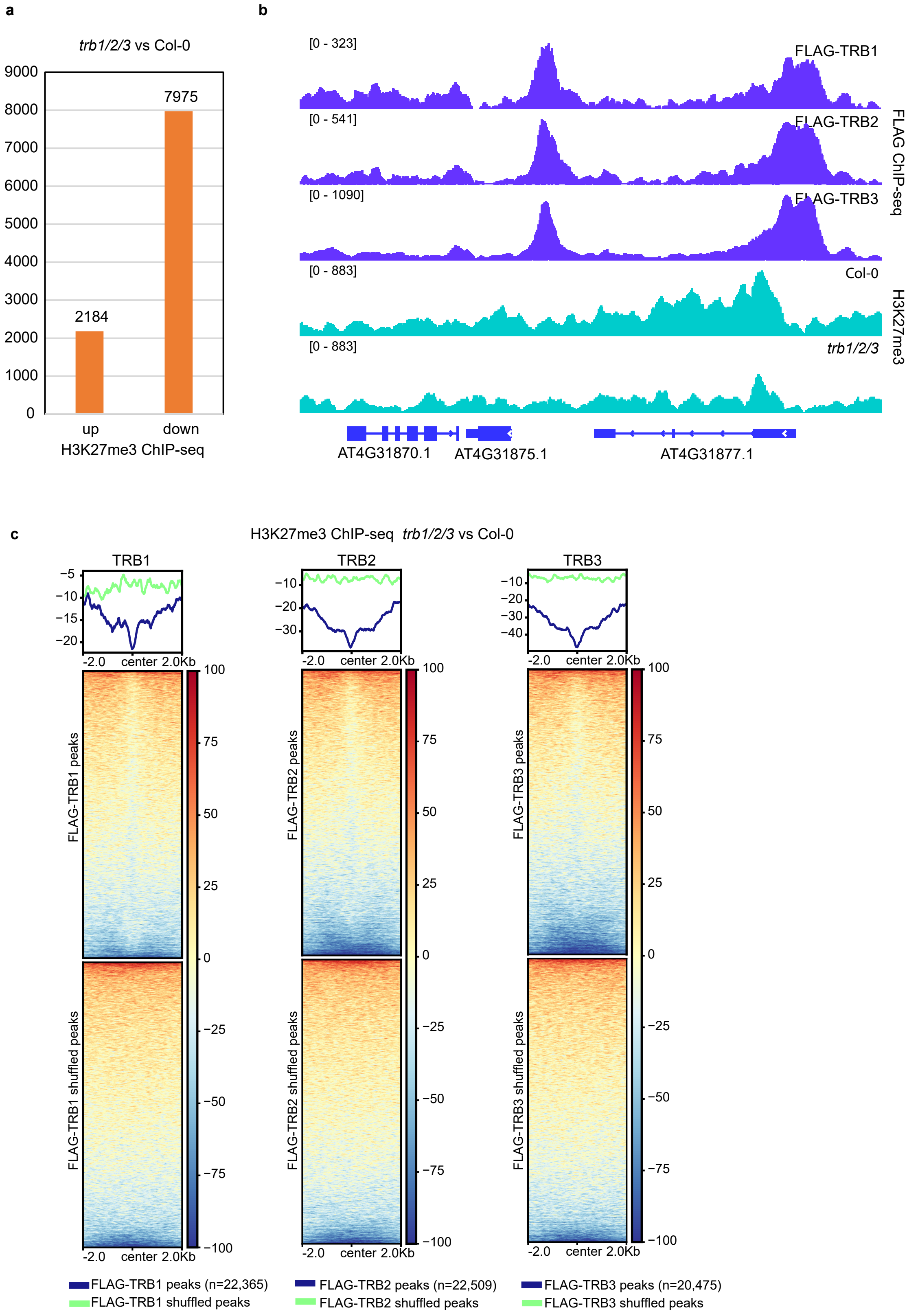


**Supplementary Fig. 6 H3K27me3 ChIP-seq signals are reduced in the *trb1/2/3* mutant. a** Bar chart indicates the number of regions with up- or down-regulated H3K27me3 ChIP-seq signals in the *trb1/2/3* mutant, respectively. **b** Screenshots showing FLAG ChIP-seq signals in FLAG-TRB1, FLAG-TRB2, and FLAG-TRB3 (top three lanes), and H3K27me3 ChIP-seq signals in Col-0 and the *trb1/2/3* mutant, over a representative region. **c** Metaplots and heatmaps representing H3K27me3 ChIP-seq signals in the *trb1/2/3* mutant versus Col-0 wild type over FLAG-TRB1 peaks and shuffled peaks (n=22,365, left panel), FLAG-TRB2 peaks and shuffled peaks (n=22,509, middle panel), and FLAG-TRB3 peaks and shuffled peaks (n=20,475, right panel).

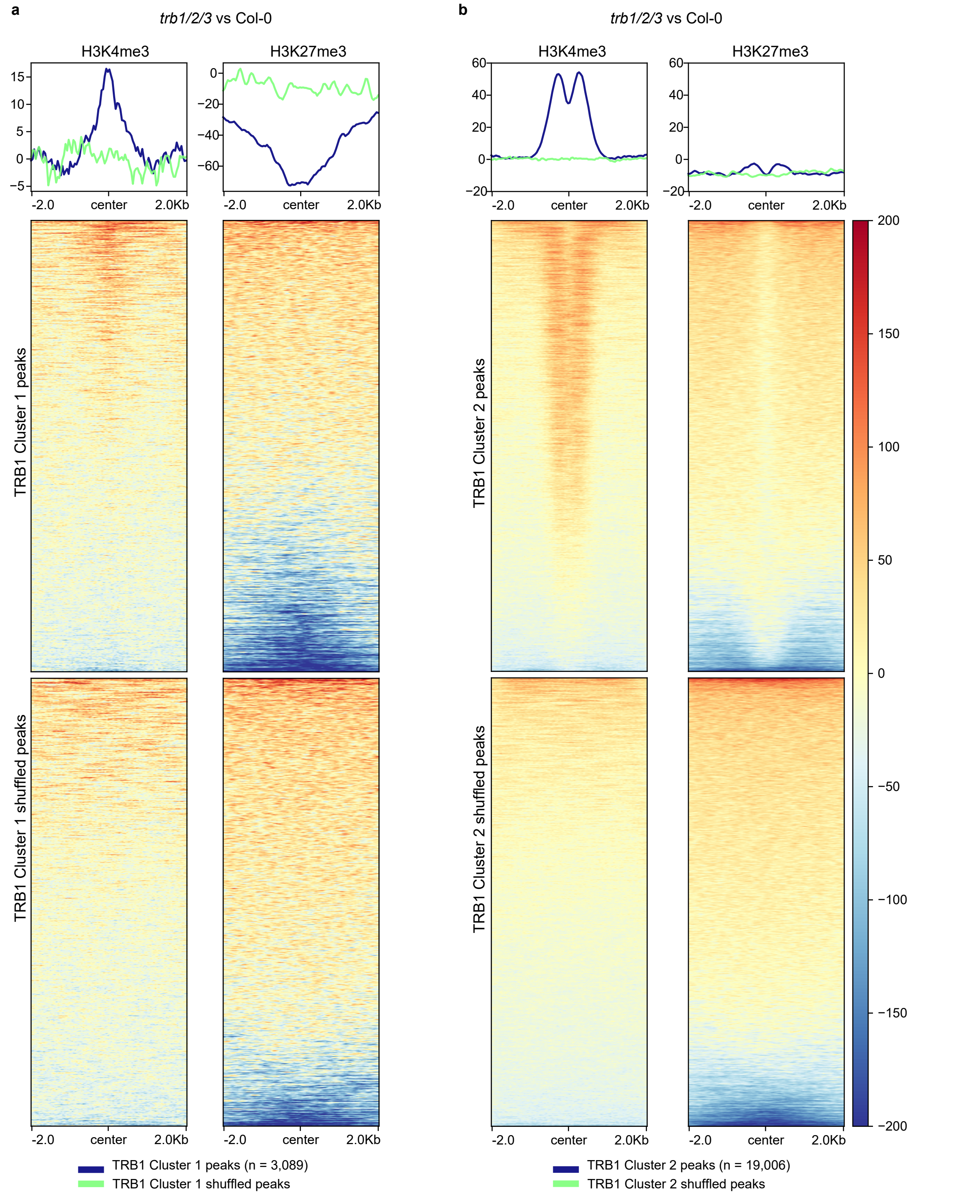


**Supplementary Fig. 7 Changes of H3K4me3 and H3K27me3 ChIP-seq signals in the *trb1/2/3* mutant displayed opposite trends at TRB1-JMJ14 co-bound regions. a-b** Metaplots and heatmaps depicting the normalized H3K4me3 and H3K27me3 ChIP-seq signals in the *trb1/2/3* mutant versus Col-0 at TRB1 Cluster 1 peaks and shuffled peaks (**a**, n=3,089), and TRB1 Cluster 2 peaks and shuffled peaks (**b**, n=19,006), respectively.

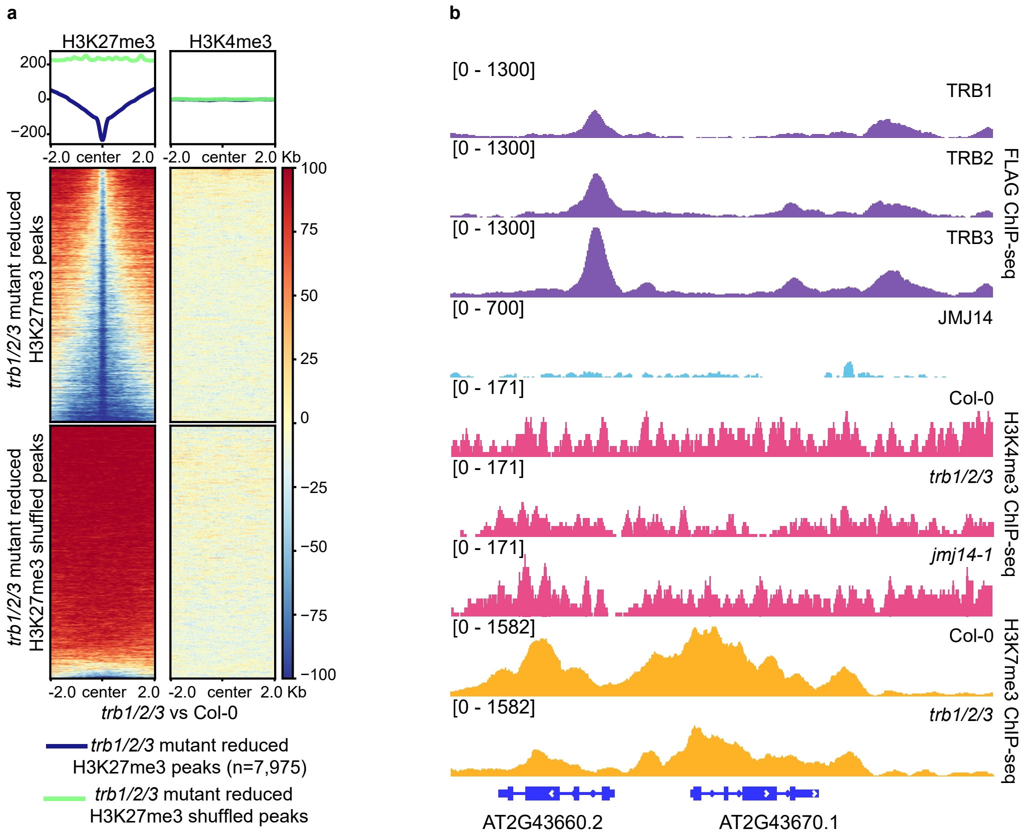


**Supplementary Fig. 8 The reduction of H3K27me3 does not always trigger an increase of H3K4me3 in the *trb1/2/3* mutant. a** Metaplots and heatmaps depicting H3K27me3 and H3K4me3 ChIP-seq signals over the reduced H3K27me3 peaks and shuffled peaks (n=7,975) in the *trb1/2/3* mutants versus Col-0 wild type. **b** Screenshots showing FLAG ChIP-seq signals of FLAG-TRB1, FLAG-TRB2, FLAG-TRB3, and FLAG-JMJ14, H3K4me3 ChIP-seq signals in Col-0, *trb1/2/3*, and *jmj14-1* mutants, and H3K27me3 ChIP-seq signals in Col-0 and the *trb1/2/3* mutant over a representative region.

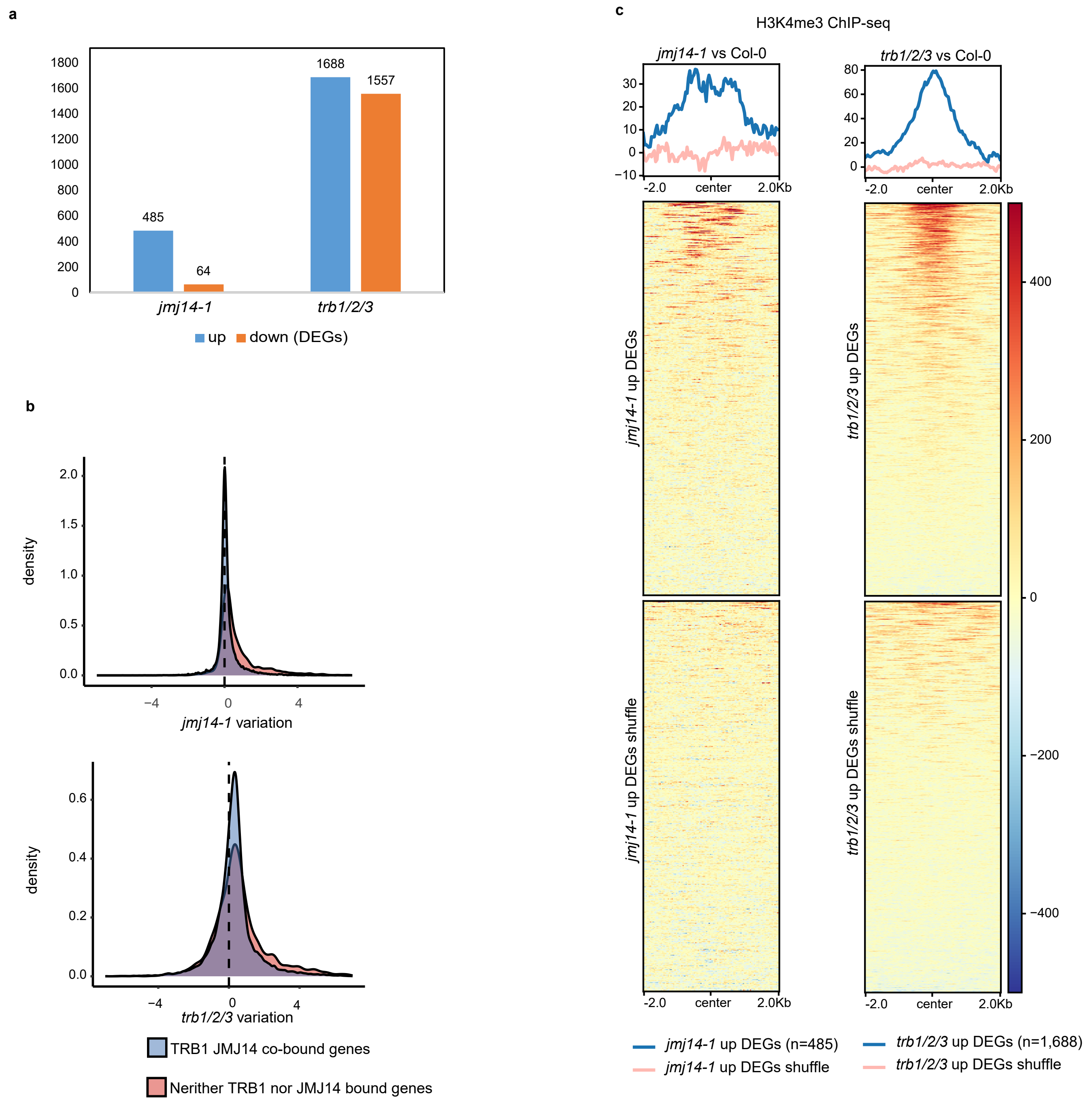


**Supplementary Fig. 9 *trb1/2/3* and *jmj14-1* up-regulated DEGs are associated with H3K4me3 induction. a** Bar chart indicates the number of up- and down-regulated DEGs in *jmj14-1* and *trb1/2/3* mutants, respectively. **b** Density plots of DEGs in *jmj14-1* (upper panel) and *trb1/2/3* (lower panel) mutants over the TRB1 and JMJ14 co-bound genes and neither TRB1 nor JMJ14 bound genes, respectively. **c** Metaplots and heatmaps representing the normalized H3K4me3 ChIP-seq signals in *jmj14-1* mutant versus Col-0 over *jmj14-1* mutant up-regulated DEGs or shuffle sites (n=485, left panel), and in *trb1/2/3* mutants versus Col-0 over *trb1/2/3* mutant up-regulated DEGs or shuffle sites (n=1,688, right panel).

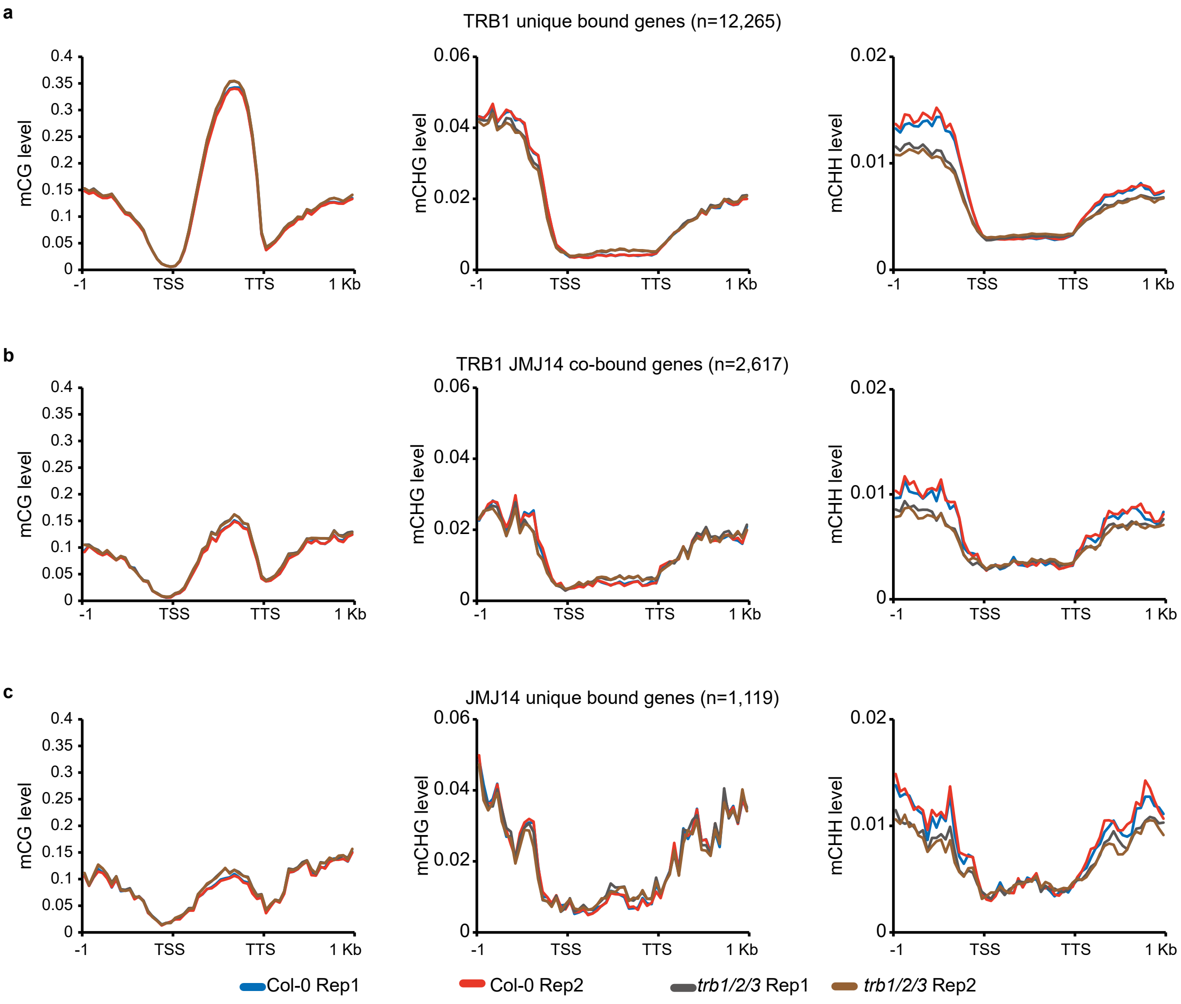


**Supplementary Fig. 10 CHH DNA methylation level is reduced in the *trb1/2/3* mutant. a-c** CG, CHG, and CHH DNA methylation levels in two replicates of Col-0 and *trb1/2/3* triple mutants over 1 kb flanks of TRB1 unique bound genes (**a**, n=12,265), TRB1-JMJ14 co-bound genes (**b**, n=2,617), and JMJ14 unique bound genes (**c**, n=1,119). The DNA methylation levels were measured by whole genome bisulfite sequencing.

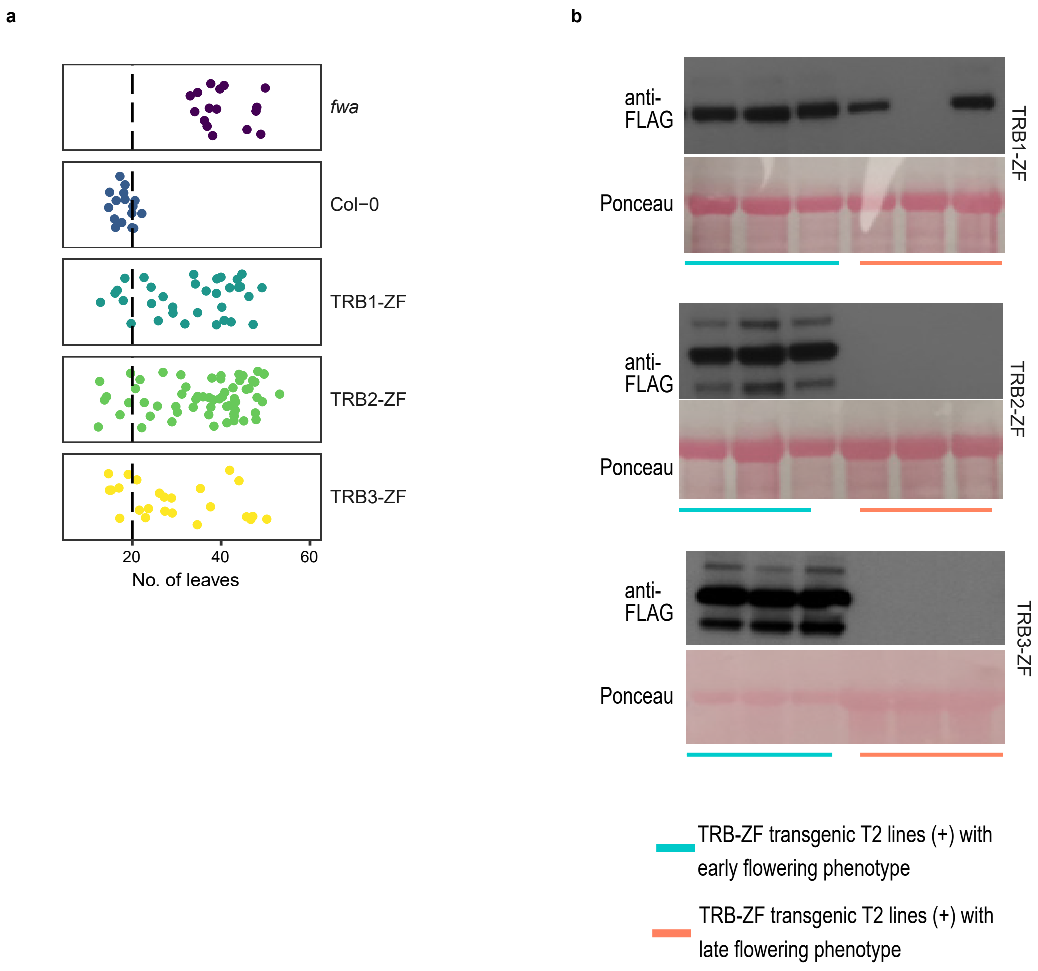


**Supplementary Fig. 11 TRB-ZF T1 lines restore an early flowering phenotype in *fwa*. a** Dot plots showing the leaf count of *fwa*, Col-0, and the T1 lines of TRB1-ZF, TRB2-ZF, and TRB3-ZF. **b** Western blot of the TRB-ZF fusions in T2 transgenic lines showing early flowering phenotypes (left three samples) and late flowering phenotypes (right three samples), respectively.

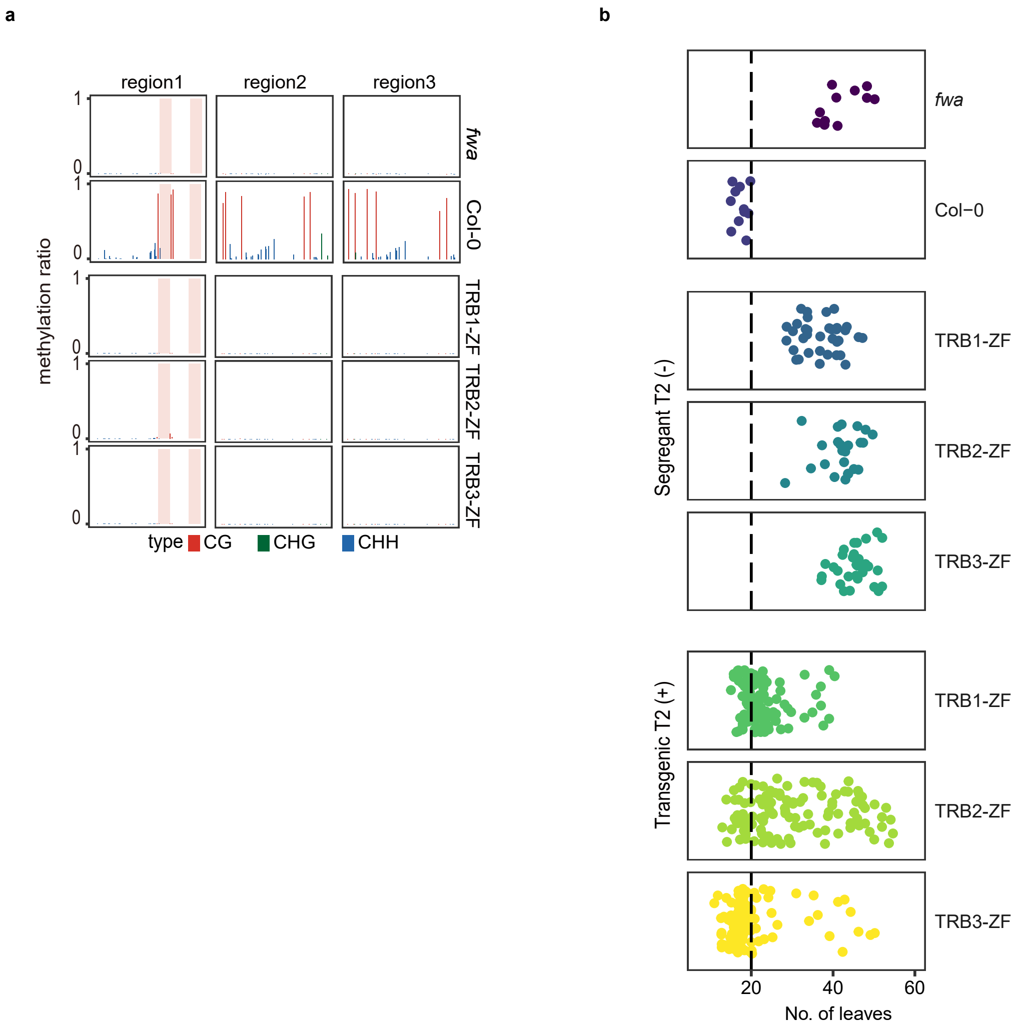


**Supplementary Fig. 12 The TRB-ZF fusions triggered early flowering phenotype is not associated with DNA methylation and is not heritable in T2 null segregant lines. a** CG, CHG, and CHH DNA methylation level over *FWA* promoter regions measured by bisulfite PCR-seq. Pink vertical boxes indicate ZF binding sites. **b** Dot plots represent the leaf counts of *fwa*, Col-0, null segregant and transgenic T2 lines of TRB1-ZF, TRB2-ZF, and TRB3-ZF, respectively.

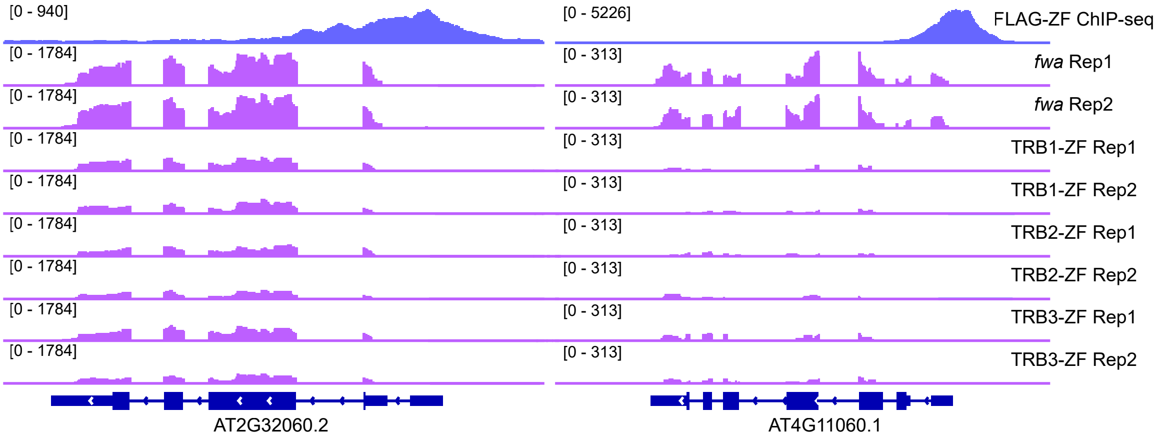


**Supplementary Fig. 13 TRB-ZF fusions triggered gene silencing over two representative ZF off-target sites.** Screenshots showing two replicates of RNA-seq signals in *fwa*, TRB1-ZF, TRB2-ZF, and TRB3-ZF over two representative ZF off target sites. FLAG-ZF ChIP-seq indicates the ZF binding sites.

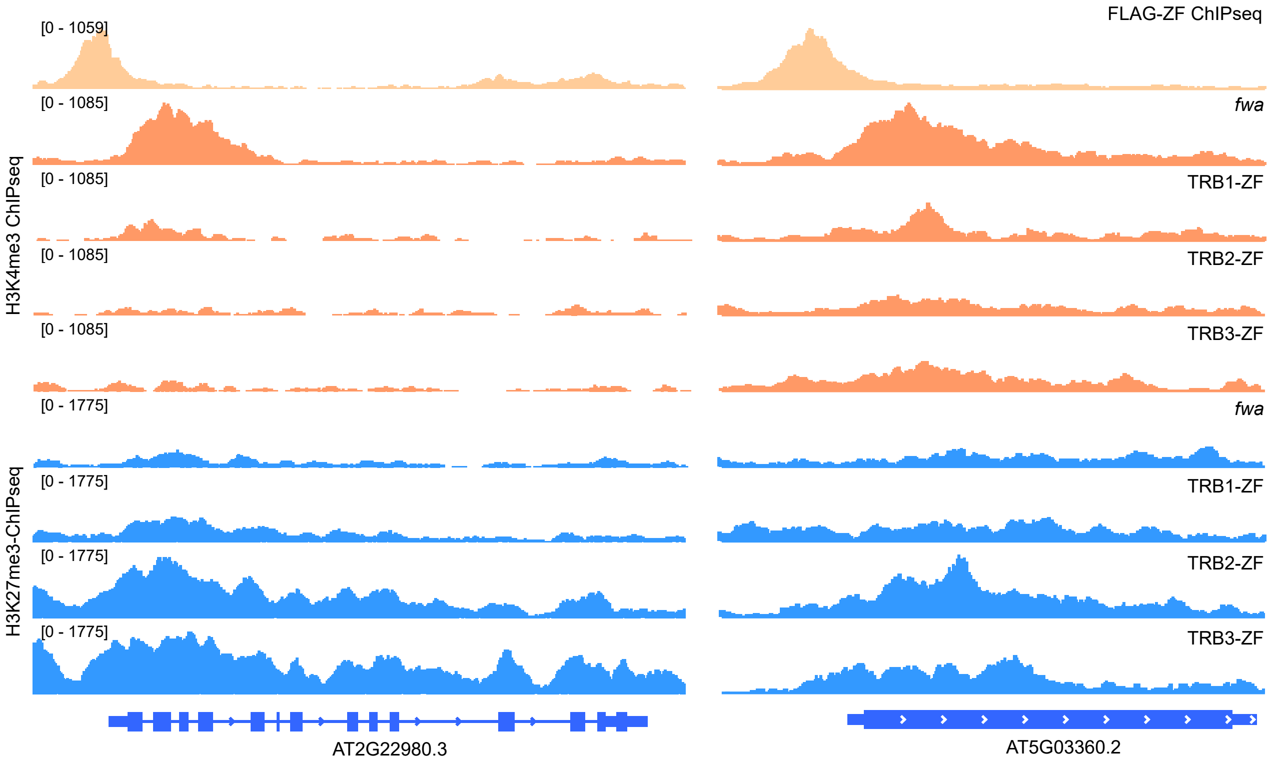


**Supplementary Fig. 14 TRB-ZF fusions triggered H3K4me3 demethylation and H3K27me3 deposition over two representative ZF off-target sites.** Screenshots of H3K4me3 and H3K27me3 ChIP-seq signals in *fwa*, TRB1-ZF, TRB2-ZF, and TRB3-ZF over two representative ZF off target sites. FLAG-ZF ChIP-seq indicates the ZF binding sites.

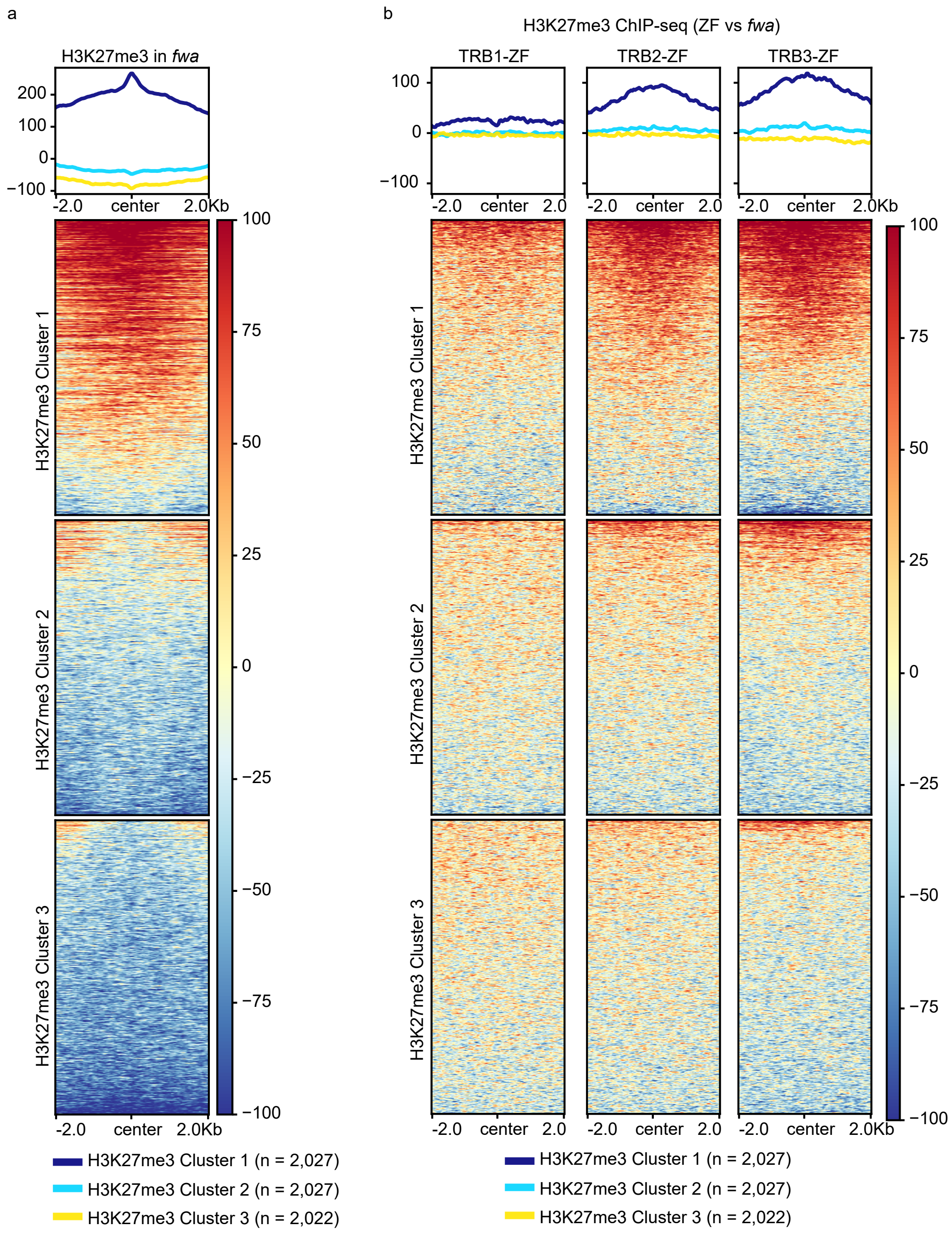


**Supplementary Fig. 15 TRB-ZFs triggered H3K27me3 deposition mainly occurs at ZF off-target sites with high levels of pre-existing H3K27me3. a** Metaplot and heatmaps representing the normalized H3K27me3 ChIP-seq signals in *fwa* plant over three clusters of ZF off-target sites with high (H3K27me3 Cluster1, n=2,027,), medium (H3K27me3 Cluster2, n=2,027), or low (H3K27me3 Cluster3, n=2,022) levels of pre-existing H3K27me3. **b** Metaplots and heatmaps showing the normalized H3K27me3 ChIP-seq signals in TRB-ZFs versus *fwa* over three H3K27me3 clusters.

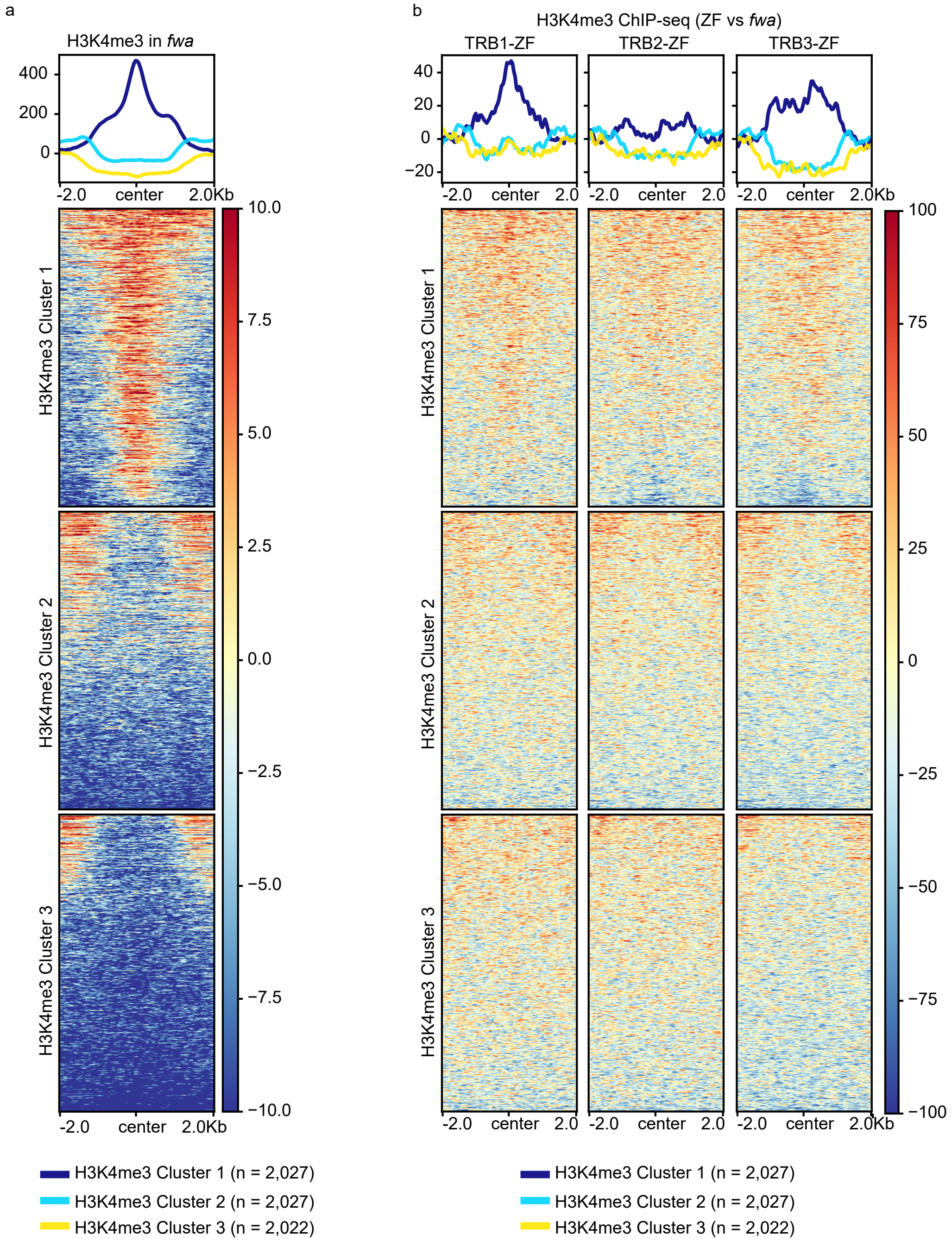


**Supplementary Fig. 16 TRB-ZFs triggered H3K4me3 removal mainly occurs at ZF off-target sites with medium or low levels of pre-existing H3K4me3. a** Metaplots and heatmaps showing the normalized H3K4me3 ChIP-seq signals in *fwa* over three clusters of ZF off-target sites with high (H3K4me3 Cluster1, n=2,027), medium (H3K4me3 Cluster2, n=2,027), or low (H3K4me3 Cluster3, n=2,022) levels of pre-existing H3K4me3. **b** Metaplots and heatmaps showing the normalized H3K4me3 ChIP-seq signals in TRB-ZFs versus *fwa* over the three H3K4me3 clusters.

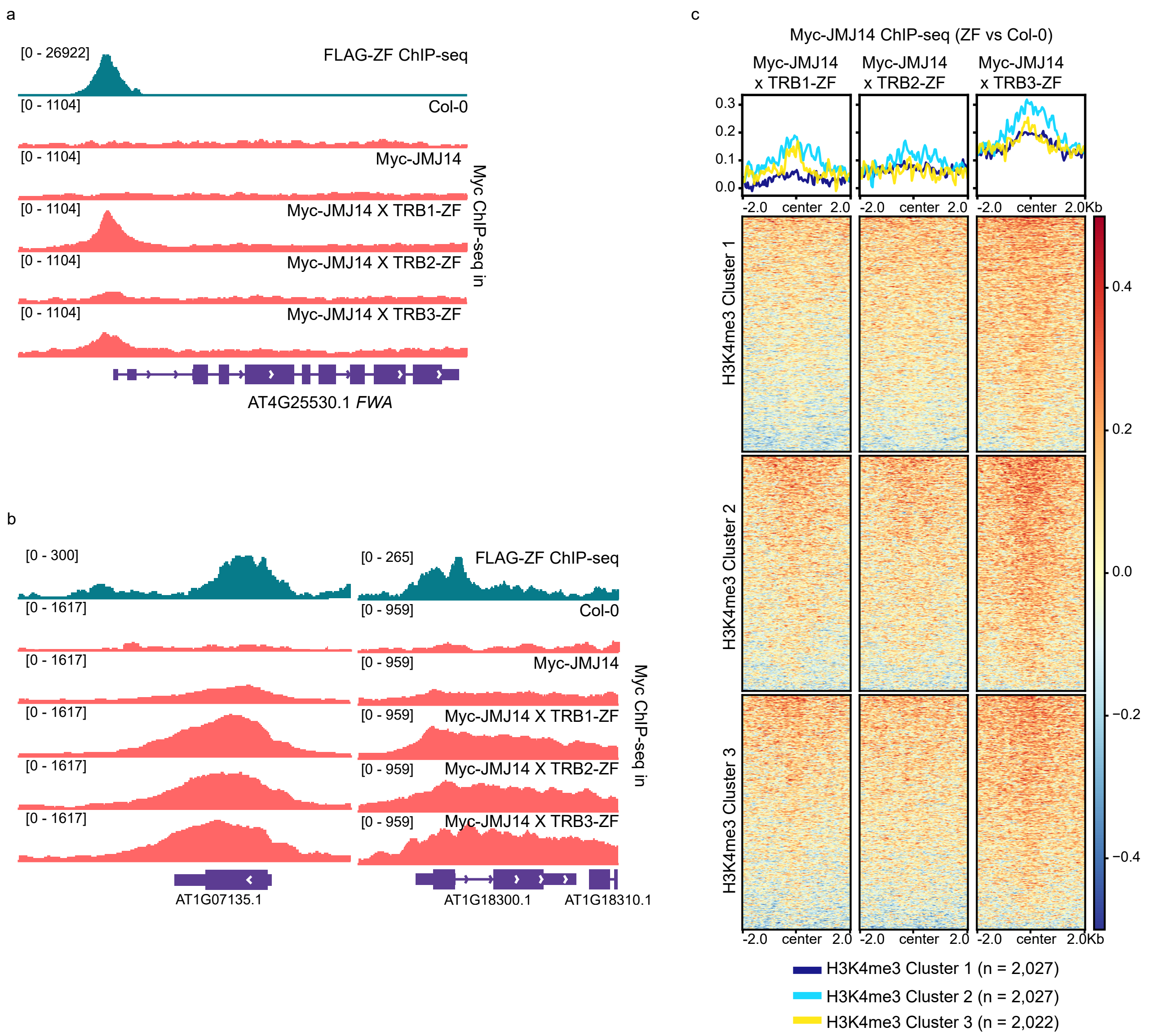


**Supplementary Fig. 17 The recruitment of JMJ14 by TRB-ZFs mainly occurs at the ZF off-target sites with medium or low levels of pre-existing H3K4me3. a-b** Screenshots of Myc ChIP-seq in Col-0 plant, Myc-JMJ14 transgenic lines in Col-0, TRB1-ZF, TRB2-ZF, and TRB3-ZF backgrounds at *FWA* (**a**) and two representative ZF off-target genes (**b**). FLAG-ZF ChIP-seq indicates the ZF binding site. **c** Normalized Myc-JMJ14 ChIP-seq signals over three H3K4me3 Clusters of ZF off-target sites in Myc-JMJ14 x TRB-ZFs crossed lines versus Myc-JMJ14 transgenic line in Col-0 background.

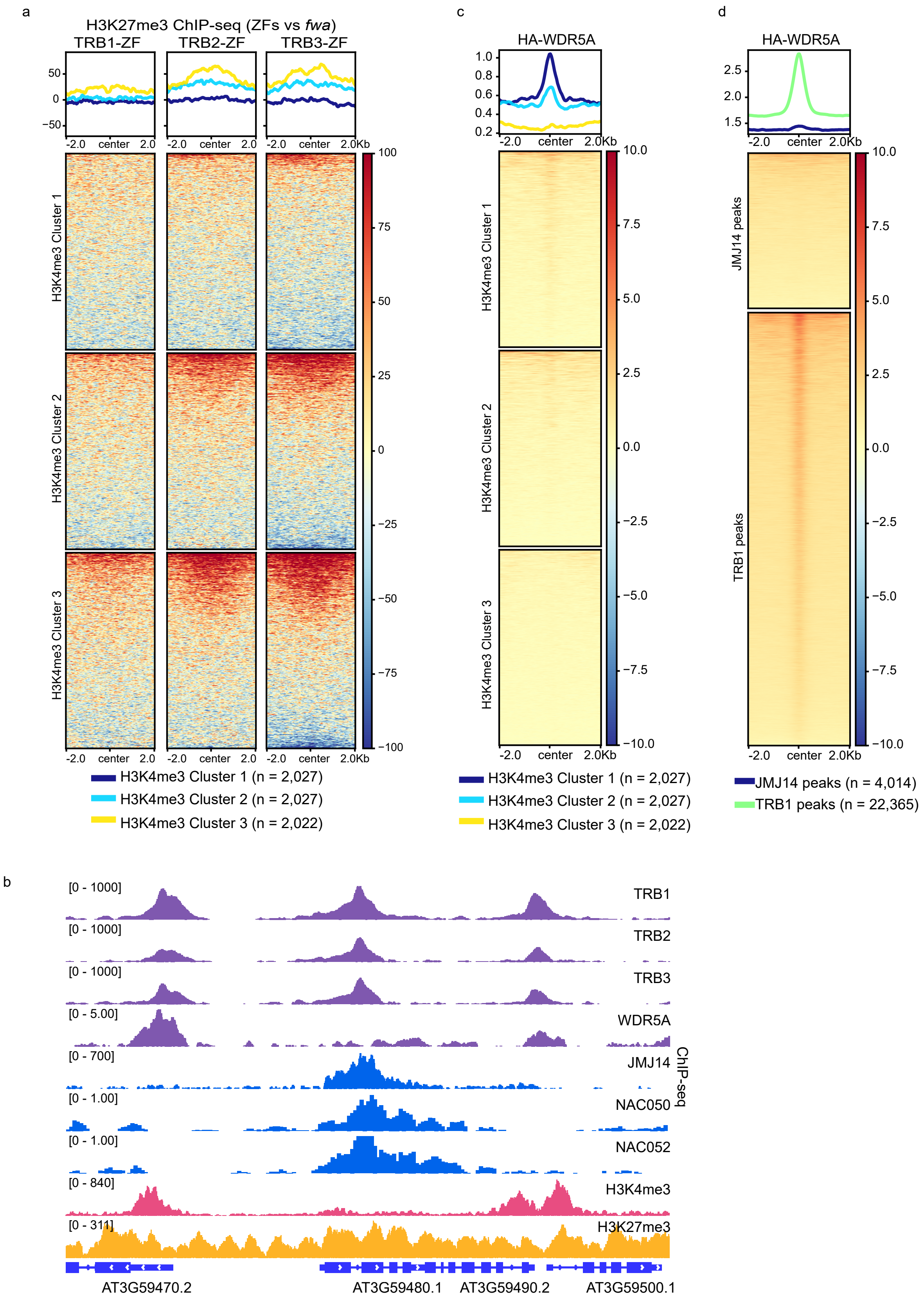


**Supplementary Fig. 18 TRB-ZF might recruit WDR5A to add H3K4me3 to ZF off-target sites with high levels of pre-existing H3K4me3. a** H3K27me3 ChIP-seq in TRB-ZFs versus *fwa* over three H3K4me3 clusters of ZF off-target sites. **b** Screenshots representing the ChIP-seq of FLAG-TRB1, FLAG-TRB2, FLAG-TRB3, FLAG-JMJ14, HA-WDR5A, HA-NAC050, HA-NAC052, H3K4me3, and H3K27me3 over a representative locus. **c-d** Metaplots and heatmaps depicting HA-WDR5A ChIP-seq signals over three H3K4me3 clusters of ZF off-target sites (**c**), or over JMJ14 peaks (n=4,014) or TRB1 peaks (n=22,36) (**d**).
